# Developmental origin and morphogenesis of the diaphragm, an essential mammalian muscle

**DOI:** 10.1101/278507

**Authors:** Elizabeth M Sefton, Mirialys Gallardo, Gabrielle Kardon

**Affiliations:** Department of Human Genetics University of Utah Salt Lake City, UT 84112

**Keywords:** diaphragm, Congenital Diaphragmatic Hernia, CDH, muscle, somite

## Abstract

The diaphragm is a mammalian skeletal muscle essential for respiration and for separating the thoracic and abdominal cavities. Development of the diaphragm requires the coordinated development of muscle, muscle connective tissue, tendon, nerves, and vasculature that derive from different embryonic sources. However, defects in diaphragm development are common and the cause of an often deadly birth defect, Congenital Diaphragmatic Hernia (CDH). Here we comprehensively describe the normal developmental origin and complex spatial-temporal relationship between the different developing tissues to form a functional diaphragm using a developmental series of mouse embryos genetically and immunofluorescently labeled and analyzed in whole mount. We find that the earliest developmental events are the emigration of muscle progenitors from cervical somites followed by the projection of phrenic nerve axons from the cervical neural tube. Muscle progenitors and phrenic nerve target the pleuroperitoneal folds (PPFs), transient pyramidal-shaped structures that form between the thoracic and abdominal cavities. Subsequently, the PPFs expand across the surface of the liver to give rise to the muscle connective tissue and central tendon, and the leading edge of their expansion precedes muscle morphogenesis, formation of the vascular network, and outgrowth and branching of the phrenic nerve. Thus development and morphogenesis of the PPFs is critical for diaphragm formation. In addition, our data indicate that the earliest events in diaphragm development are critical for the etiology of CDH and instrumental to the evolution of the diaphragm. CDH initiates prior to E12.5 in mouse and suggests that defects in the early PPF formation or their ability to recruit muscle are an important source of CDH. Also, the recruitment of muscle progenitors from cervical somites to the nascent PPFs is uniquely mammalian and a key developmental innovation essential for the evolution of the muscularized diaphragm.

**Highlights:**

- Diaphragm development begins with emigration of muscle progenitors from cervical somites.
- Phrenic nerve axons follow muscle path towards nascent pleuroperitoneal folds (PPFs).
- PPFs are the target of muscle migration and phrenic nerve axon projection
- PPF expansion precedes and likely directs muscle, nerve, and vasculature development.
- Early defects in PPFs and muscle recruitment are likely a source of CDH.

## Introduction

The diaphragm is an essential mammalian skeletal muscle that has two critical functions (Merrell and Kardon, 2013). As a domed muscle lying at the base of the thoracic cavity, its contraction expands the thoracic cavity and drives the inspiration phase of respiration (Campbell et al., 1970). In addition, as a muscle separating the thoracic and abdominal cavities, it serves as an important barrier between these two regions (Perry et al., 2010). Hence morphogenesis of a fully functional diaphragm is critical for mammalian respiration and for proper organization of the internal viscera. The essential barrier function of the diaphragm is disrupted in the common birth defect, Congenital Diaphragmatic Hernia (CDH). In CDH, diaphragm development is defective, leading to a weak or incompletely formed diaphragm, and results in 50% neonatal mortality and long-term morbidity (Pober, 2007). While recent genetic and developmental studies in mice and humans (Kardon et al., 2017) give us insights into the mechanisms regulating diaphragm development and the etiology of CDH, critical gaps in our knowledge remain.

The respiratory and barrier functions of the diaphragm are primarily carried out by the costal diaphragm (Fig. 1). The costal diaphragm consists of a radial array of myofibers, each surrounded by connective tissue, which extend from the ribs to the central tendon. These myofibers are innervated by bilateral phrenic motor nerves. Originating from the cervical region (C3-C5), the axons of the right and left phrenic nerves descend along the vertebrae and pericardium, pierce the diaphragm, and subdivide into three branches innervating the ventral and dorsal costal muscle as well as the crural muscle (which surrounds the esophagus and is important for swallowing) (Allan and Greer, 1997). All the tissues of the diaphragm are vascularized by three sets of arteries: phrenic, internal thoracic, and intercostal arteries (Stuelsatz et al., 2012).

**Figure 1:**
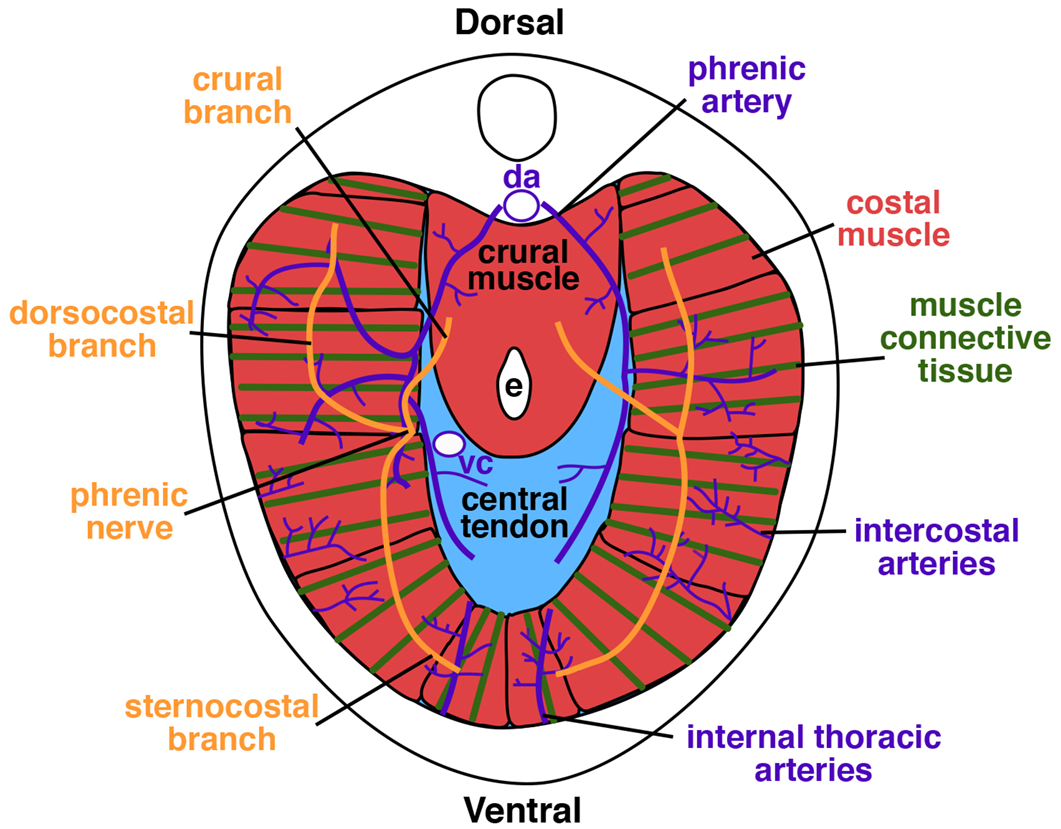
Anatomy of adult diaphragm, cranial view.

The adult diaphragm derives from multiple embryonic sources. Specific to the developing diaphragm are the pleuroperitoneal folds (PPFs; reviewed in Kardon et al., 2017; Merrell and Kardon, 2013). The PPFs are two transient pyramidal-shaped structures located between the thoracic (pleural) and abdominal (peritoneal) cavities (Babiuk et al., 2003; Merrell et al., 2015). The PPFs expand dorsally and ventrally across the septum transversum, a thin membrane on the cranial surface of the liver, and give rise to the diaphragm’s muscle connective tissue and central tendon (Merrell et al., 2015). There is some evidence that the PPFs originate in the cervical region (Hirasawa et al., 2015), but nascent PPFs have never been explicitly labeled and their early development tracked. The somites are the source of the diaphragm’s muscle, and muscle progenitors emigrate from the somites into the PPFs (Allan and Greer, 1997; Babiuk et al., 2003; Dietrich et al., 1999; Merrell et al., 2015). The axons of the phrenic nerve arise from the C3-C5 segments of the neural tube, and similar to the muscle progenitors, project to the PPFs (Allan and Greer, 1997, 1998; Babiuk et al., 2003). The migration of muscle and projection of nerve to the PPFs are key developmental events, but which somites give rise to the diaphragm’s muscle and the spatial-temporal relationship of muscle migration and phrenic nerve outgrowth to the PPFs has not been determined. The final major component of the diaphragm is the vasculature, but its embryonic source is not known.

The PPFs are critical for diaphragm development (Merrell et al., 2015). By embryonic day (E) 11.5 in the mouse the muscle progenitors are present in the core of the PPFs, and the subsequent dorsal and ventral expansion (E11.5-E16.5) of the PPFs carries the muscle throughout the developing diaphragm and so regulates muscle morphogenesis (Merrell et al., 2015). The outgrowth of the three phrenic nerve axon branches lags behind the leading edge of the muscle progenitors (Allan and Greer, 1997, 1998; Babiuk et al., 2003) and so likely is also controlled by the expansion of the PPFs. Interestingly, while the PPFs give rise to the muscle connective tissue and central tendon, in the adult the muscle and phrenic nerve are excluded from the central tendon and reside only in the connective tissue region. Essential to the survival of these tissues is the vasculature, and while the adult vascular network has been described in detail (Stuelsatz et al., 2012), how it forms has not.

Defects in diaphragm development are the cause of the common (1/3,000 total births) birth defect, CDH (Torfs et al., 1992). In CDH, weaknesses in the developing diaphragm allow abdominal contents to herniate into the thoracic cavity and impede lung development. The associated lung hypoplasia is the cause of the high rates of neonatal mortality and long-term morbidity associated with CDH (Lally, 2016). CDH usually appears as an isolated defect, but in 30-40% of cases CDH patients have other developmental malformations (Pober, 2007); besides the lungs, defects in the heart are particularly common, and CDH is also associated with central nervous system and limb abnormalities (Pober, 2007). A noted feature of CDH is that hernias preferentially occur on the left side of the diaphragm (Pober, 2007), although there is currently no clear consensus on whether right or left hernias are associated with a worse clinical outcome (Azarow et al., 1997; Duess et al., 2015; Grizelj et al., 2017; Losty et al., 1998; Schaible et al., 2012; Skari et al., 2002; Skari et al., 2000).

Despite the prevalence and severity of CDH, the genetic and cellular etiology is incompletely understood (Kardon et al., 2017). The genetic etiology of CDH is heterogeneous and has been associated with aneuploidies, cytogenetic rearrangements and copy number variants involving multiple chromosomes as well as mutations in many individual genes (Kardon et al., 2017; Pober, 2007). Of the greater than 100 genes that have been implicated by human genetic studies, fewer than 20 have been verified as causing CDH via mouse genetic studies (Kardon et al., 2017). Several studies conditionally deleting CDH genes in specific embryonic tissues in mice have definitively shown that mutations in the PPFs (or the mesothelium associated with the PPFs) are a source of CDH (Carmona et al., 2016; Merrell et al., 2015; Paris et al., 2015).

Mutations in in the PPFs lead to hernias through a variety mechanisms: decreased proliferation, increased apoptosis, and changes in differentiation of PPF cells; non cell-autonomous effects on muscle progenitor proliferation and apoptosis; and changes in diaphragm morphogenesis (Carmona et al., 2016; Merrell et al., 2015; Paris et al., 2015). Surprisingly, these studies demonstrated that diaphragm defects are present (Carmona et al., 2016; Merrell et al., 2015) or genes are required (Paris et al., 2015) prior to E12.5. While normal diaphragm morphogenesis of PPFs and muscle at E12.5 – E16.5 has been carefully analyzed (Babiuk et al., 2003; Merrell et al., 2015), the earlier developmental events have not been similarly analyzed.

Here we address the lack of a comprehensive analysis of the developmental origin and coordinated development of multiple tissues during diaphragm morphogenesis that result in the formation of a functional diaphragm. We use a developmental series of mouse embryos in which somite-derived muscle, neural tube-derived phrenic nerve, PPF-derived muscle connective tissue and central tendon, and vasculature are either genetically or immunofluorescently labeled and analyzed in whole mount. Our four-dimensional reconstruction of the developing diaphragm uniquely elucidates the formation of the PPFs, origin and early migration of muscle progenitors, phrenic nerve outgrowth, and coordinate morphogenesis of the PPFs, muscle, nerve, and vasculature. In addition, the spatial-temporal relationship between the developing diaphragm and the surrounding heart and its associated vessels, lungs, and liver is described. Our work provides important insights into the normal development of the diaphragm, the etiology of CDH, and the evolutionary origins of the mammalian diaphragm.

## Materials and Methods

### Mice and staging

All mice have been previously published. We used Prx1Cre^Tg^ (Logan et al., 2002), *Pax3^Cre^* (Engleka et al., 2005) Cre alleles and *Rosa26^LacZ^* (Soriano, 1999) and *Rosa26^mTmG^* (Muzumdar et al., 2007) Cre-responsive reporter alleles. Mice were back-crossed on to the C57/BL6J background. Animal experiments were performed in accordance with protocols approved by the Institutional Animal Care and Use Committee at the University of Utah.

Embryos were staged as E0.5 on the day dams had a vaginal plug. To more finely stage embryos, the total number of somites were counted. In addition, to identify particular cervical somites we used the system of Spörle and Schughart (1997). Prior to formation of dorsal root ganglia, the characteristic “triad” of the last occipital somite, first and second cervical somites were uniquely identifiable. Later in development, the rectangular first cervical spinal ganglia allowed for identification of the first cervical somite. Five occipital somites were presumed, although the anterior-most occipital somites may disperse early in development (O’Rahilly and Müller, 1984).

### Wholemount immunohistochemistry, immunofluorescence, and in situ hybridization

For whole mount immunofluorescence, embryos were fixed overnight in 4% paraformaldehyde in PBS at 4°C, bleached overnight in Dent’s bleach (1:2 30% hydrogen peroxide: Dent’s fix) at 4°C, then stored in Dent’s fix (1:4 DMSO: methanol) at 4°C for at least five days. Embryos were washed, blocked in 5% goat serum with 20% DMSO, and then incubated in primary and secondary antibodies for 48 hours each at room temperature. For CD31, embryos were blocked in 10% goat serum, 0.5% Triton X-100 and 2% BSA. Primary CD31 and secondary antibodies were used with 5% goat serum, 0.2% triton X-100 and 2% BSA. Embryos were then washed in PBS and methanol and cleared using BABB (1 Benzyl Alcohol: 2 Benzyl Benzoate). For detection of HRP-conjugated secondary antibodies, embryos were incubated in 10 mg Diaminobenzidine tetrahydrochloride in 50 ml PBS with 7ul 30% hydrogen peroxide for 20 minutes.

For a list of antibodies used, see Supplemental Table 1.

Whole mount in situ hybridization was performed according to the protocol in (Riddle et al., 1993)

### Microscopy and three-dimensional rendering

Fluorescence images were taken either on a Nikon A1 or a Leica SP8 microscope. Optical stacks of whole-mount images were rendered and structures highlighted using Fluorender (Wan et al., 2009).

## Results

### The diaphragm originates at cervical levels and then “descends” to the base of the thorax

Although the adult diaphragm resides at the base of the thorax, the early nascent diaphragm forms at the cervical level, adjacent to somites C3-C5. To track the location and timing of the caudal descent of the diaphragm, we labeled the nerves via immunohistochemistry in whole mount. The axons of the phrenic nerves extend to the developing diaphragm region, and so the distal-most extent of the phrenic nerve axons approximates the location of the nascent diaphragm (see below). At E11.25 (embryo has a total of 42 somites), the axons of the phrenic nerve are first visible, extend primarily medially, and indicate that the nascent diaphragm is at the level of C6 (Fig. 2A-B). As indicated by the distal extent of the phrenic, by E11.5 (45 total somites) the diaphragm has descended to C8 and by E12.5 to T5 (Fig. 2C-F). The diaphragm reaches its final position at the base of the thorax by E13.5 (data not shown and Greer et al., 1999). As is visible in whole mount, the descent of the diaphragm is also accompanied by the caudal displacement of the heart and lungs. The diaphragm and the organs of the thoracic cavity descend from cervical levels to their final location between E11 and E13.5.

**Figure 2:**
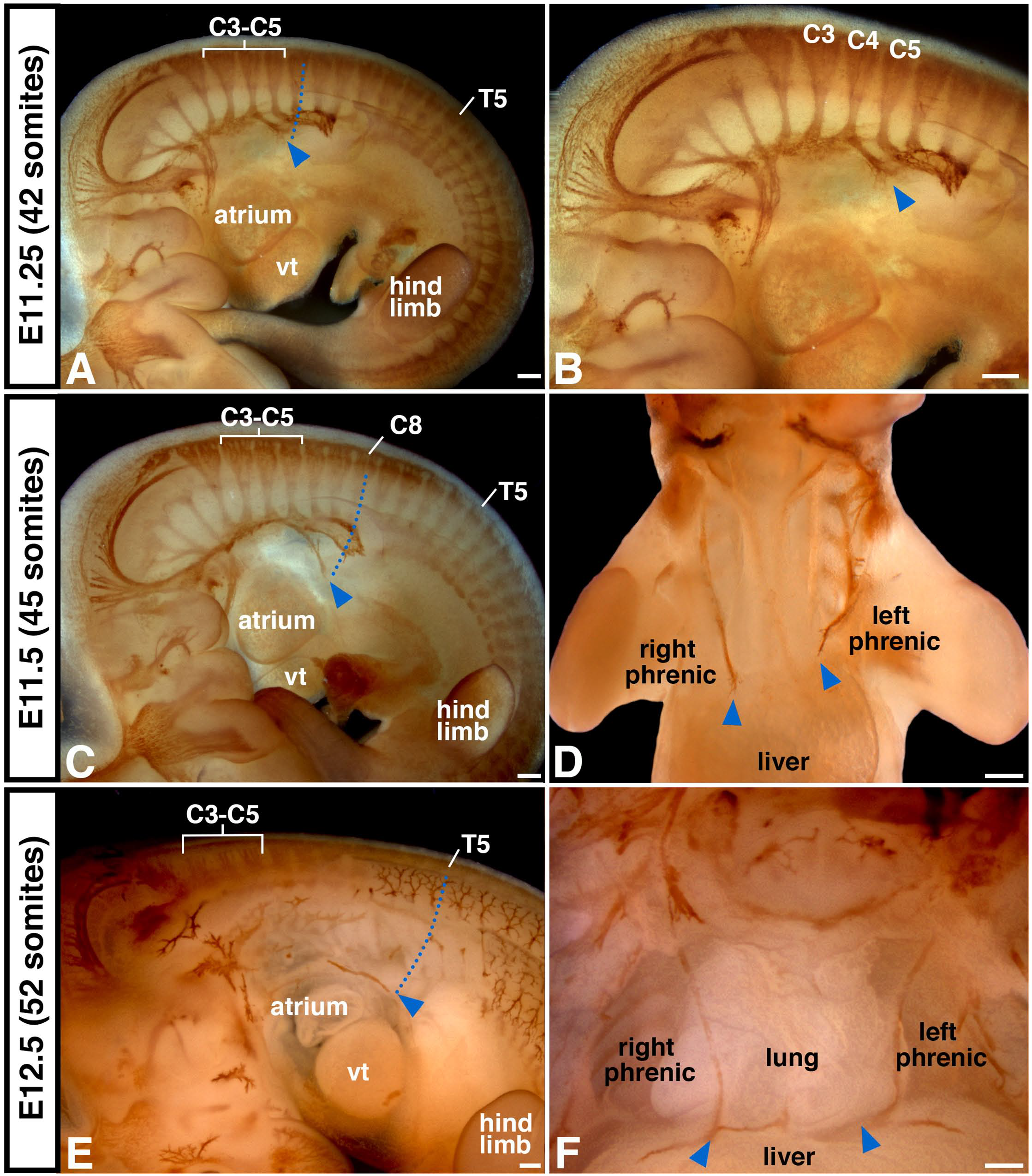
As indicated by phrenic nerve outgrowth, the diaphragm originates at the cervical level and descends to the base of the thorax. (A-B) Phrenic nerve, labeled via neurofilament antibody, emerges at E11.25 from the spinal cord at C3-C5 level and extends medially to the developing diaphragm located at C5 level (blue arrowheads and dotted line). (C-F) By E11.5 phrenic nerve axon tips and developing diaphragm (blue arrowheads) have descended to C8 level and by E12.5 to T5 level. (D) Note left phrenic nerve consistently lags behind right nerve. vt, ventricle. Scale bar A-D = 200um, E = 100um, F = 200um.

### Muscle progenitor delamination and migration precedes projection of the phrenic nerve to the nascent diaphragm region

Two important components of the diaphragm, the muscle and nerve, travel from cervical regions to their target in the nascent PPFs. We compared the spatiotemporal relationship between early muscle progenitor migration and phrenic nerve axon outgrowth using a series of embryos genetically and immunofluorescently labeled in whole mount. To determine the somitic origin of diaphragm muscle, we used *Pax3^Cre/+^; Rosa26^mTmG/+^* mice (Engleka et al., 2005; Muzumdar et al., 2007), where Cre-mediated recombination in the somite labels myogenic cells as GFP+ (Hutcheson et al., 2009). As the transcription factor Pax3 also labels neural crest cells, we verified that GFP+ cells are indeed muscle progenitors by comparing GFP and Sox10 expression (which labels neural crest; McKeown et al., 2013) and found no overlap between Sox10 and GFP in the region and time period of analysis (data not shown). In addition, we compared GFP with the expression of the receptor tyrosine kinase *Met* (which labels all migratory muscle progenitors; Bladt et al., 1995; Dietrich et al., 1999) and found that GFP and *Met* label the same population of muscle progenitors (Fig. S1). To compare muscle migration with phrenic nerve outgrowth, *Pax3^Cre/+^; Rosa26^mTmG/+^* embryos were also labeled with neurofilament immunofluorescence. In addition, we used Tomato fluorescence (which marks Cre-cells) to demarcate all other structures in the embryo.

The migration of muscle progenitors to the developing diaphragm occurs in a highly stereotyped and reproducible fashion. Beginning at E9.5 (embryos with 25 total somites), the first muscle progenitors delaminate from the cervical somites (Fig. 3A). By E10.0, a stream of muscle progenitors migrates medial to the forelimb muscle progenitors (Fig. 3B). Notably, this stream of diaphragm muscle progenitors is most closely associated with the muscle progenitors migrating to the forelimb and is distinct from the hypobranchial progenitors providing tongue and throat muscles (Fig. 3C; Lours-Calet et al., 2014). Between E10.5 and E11.5 this stream of muscle progenitors migrates towards the midline and comes to reside in a small region bounded by the lungs, heart, sinus venosus (the vessel that transmits venous blood to the atrium of the embryonic heart), and liver (Fig. 3C-IThis region corresponds to the region of the nascent PPFs (Fig. 4 and see below), and by E12.0 the muscle progenitors coalesce and are completely surrounded by well-defined PPFs (Fig. 5 and 6A). Muscle delamination and migration from the somites to the nascent PPFs occurs strictly between E9.5 and E11.5. A comparison of muscle migration on left and right sides of the embryo reveals that left and right migration are often not perfectly coordinated (data not shown), but there is no consistent trend whereby one stream migrates in advance of the other. Once the muscle progenitors are encased in the PPFs, the progenitors, PPFs, lungs, heart, sinus venosus, and liver are all translocated caudally between E11.25 and E13.5 (Fig. 3E-F, H-I; Fig. 2, and data not shown).

**Figure 3:**
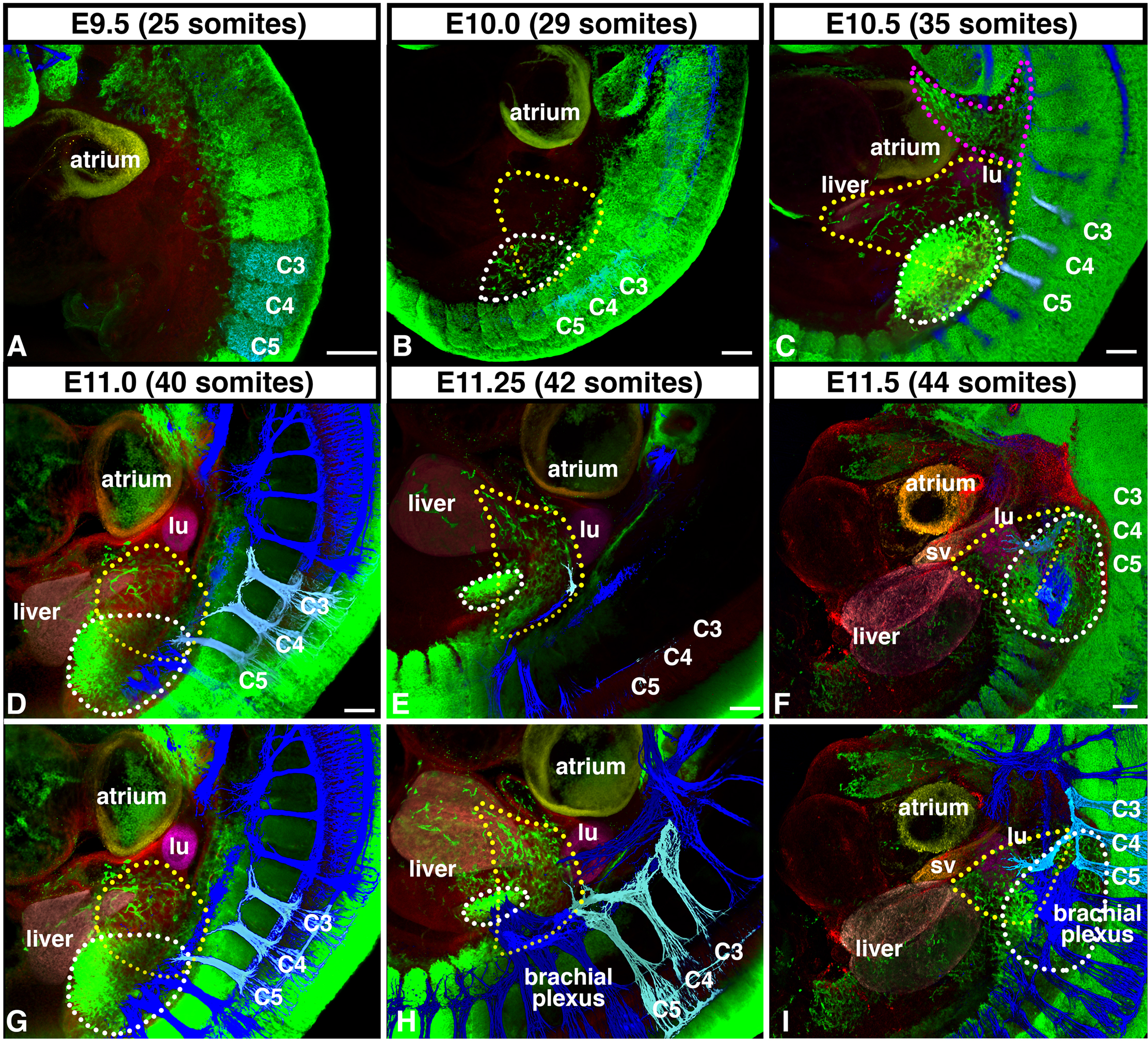
Diaphragm muscle originates from cervical somites and migrates medially to PPFs in advance of phrenic nerve. (A-B) Beginning at E9.5, diaphragm muscle progenitors (green, and outlined in white in B), labeled via *Pax3^Cre/+^;Rosa26^mTmG/+^*, delaminate from somites C3-5 (and potentially C2 and C6) and migrate medial to limb muscle progenitors (outlined in yellow in B) that delaminate from somites C4-T2 and are distinct from hypobranchial muscle stream (outlined in pink in C). (C-E) At E10.0-11.25, Diaphragm muscle progenitors migrate just caudal to lung buds (Tomato+, pseudo-colored in fuchsia) and cranial to liver lobes (Tomato+, pseudo-colored in pink). (F) By E11.5 muscle progenitors reach developing diaphragm region, wedged between the lung (fuchsia), liver (pink), and sinus venosus (Tomato+, pseudo-colored in orange). Beginning at E11.25 phrenic nerve (labeled via neurofilament immunofluorescence, pseudo-colored in pale blue) exits brachial plexus (all nerves in dark blue) and extends towards developing diaphragm, but does not reach diaphragm until E11.75 (See Fig 2). (B-I) Diaphragm muscle progenitors encircled by yellow dotted lines and limb muscle progenitors encircled by white dotted lines. (D-I) To image muscle progenitors migrating to the diaphragm, lateral part of the embryo, including most of the forelimb, was removed. (A-I) A-F imaged in depth mode, G-H identical to D-F except nerves (blue) are layered on top to reveal network of nerves. lu, lung; sv, sinus venosus. Scale bar A-F = 100 um.

**Figure 4:**
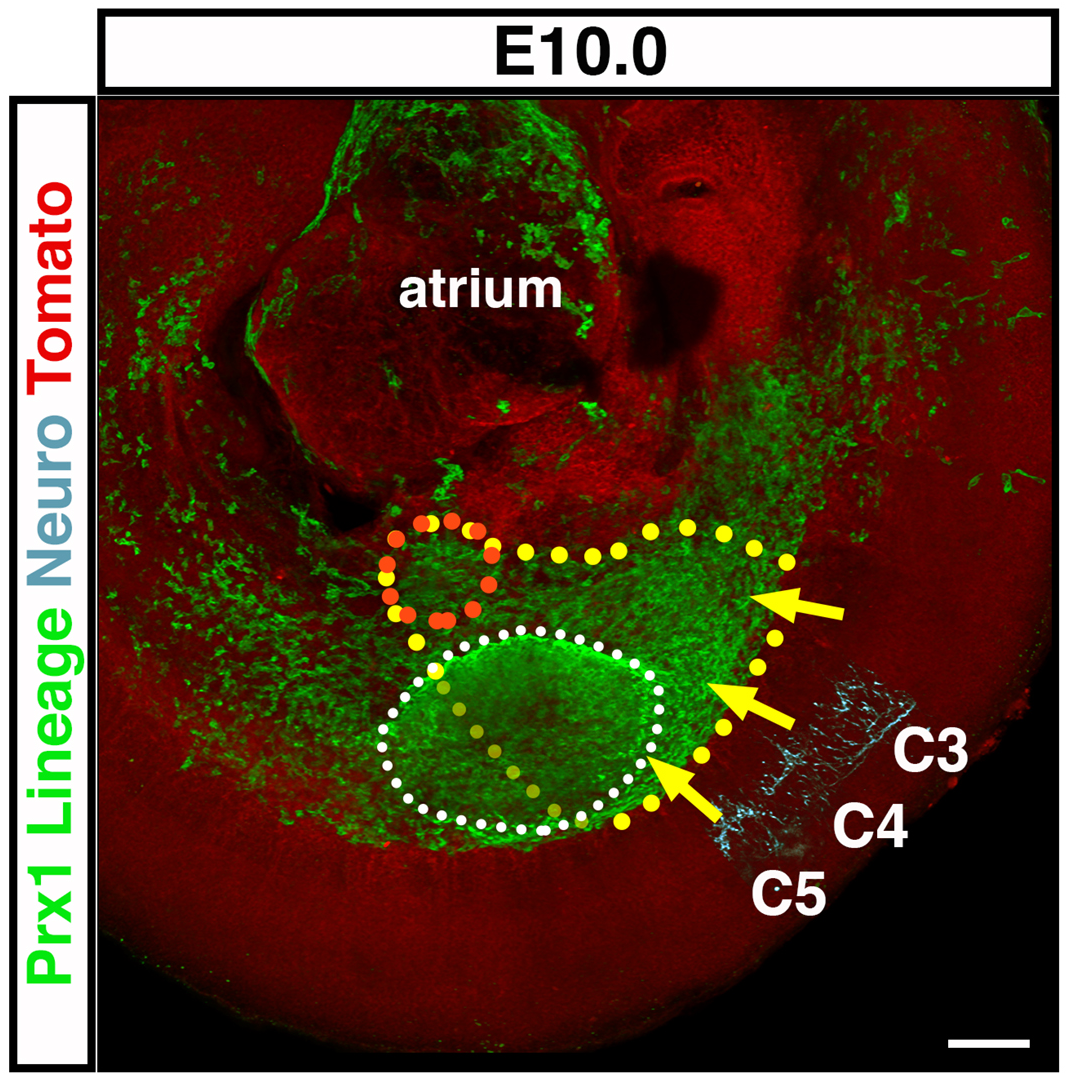
Nascent PPFs are lateral-plate derived and the target of diaphragm muscle progenitors and phrenic nerve. At E10.0 just forming PPFs are a medial extension of lateral plate, as labeled by *Prx1Cre^Tg/+^;Rosa26^mTmG/+^* (and outlined in orange) and are the target of muscle progenitors migrating from somites C3-C5 (outlined in yellow, as extrapolated from Fig 3C) and phrenic nerve axons. Forelimb is outlined in white. Scale bar = 100um.

**Figure 5:**
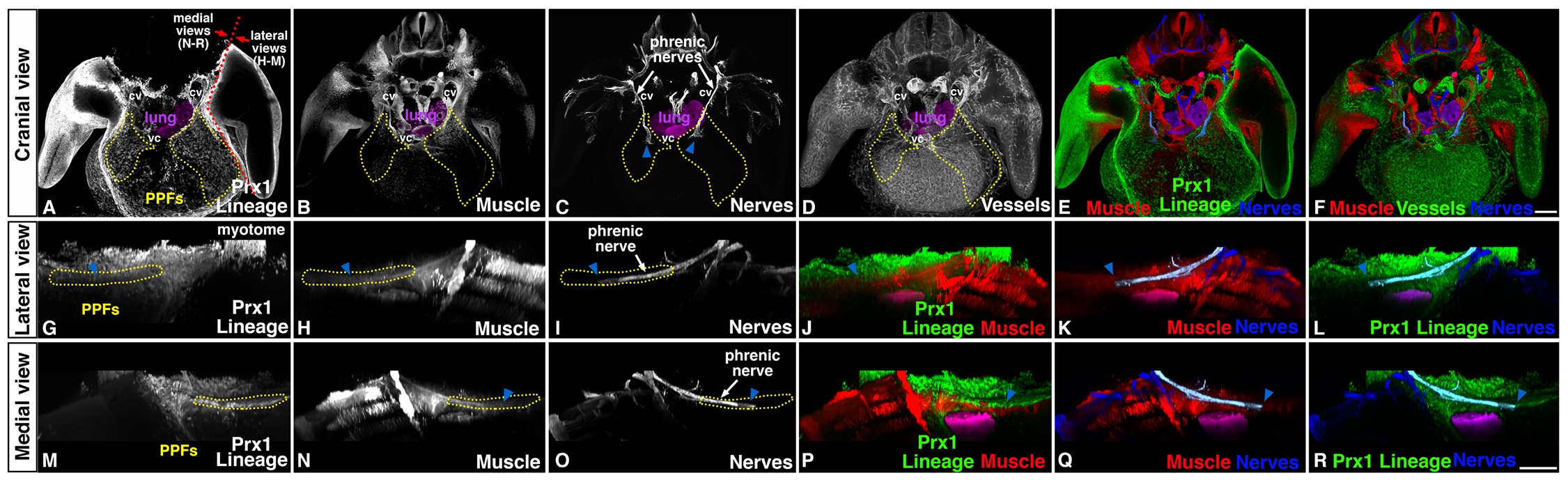
At E12.0 muscle, nerve, vasculature, and PPF-derived connective tissue and tendon progenitors are present in developing diaphragm region. (A-C, E, G-R) Into PPFs (as shown by GFP+ *Prx1*-derivatives in *Prx1Cre^Tg/+^;Rosa26^mTmG/+^*), muscle progenitors (Pax7+MyoD+ via immunofluorescence) migrate from the somites and phrenic nerve axons (Neurofilament+ via immunofluorescence) extend from the neural tube. Muscle progenitors and phrenic axon tip completely surrounded by PPFs. (D, F) Also at E12.0 vasculature (CD31+ via immunofluorescence) beginning to develop in PPF region in close association with muscle progenitors. cv, cardinal vein; vc, vena cava. Scale bar A-F = 200 um. Scale bar G-R = 200um.

While the diaphragm’s muscle progenitors begin migration at E9.5, distinct phrenic nerve axons are first visible at E11.25 (Fig. 3E, H). The phrenic nerve derives from nerve roots C3-C5, and its bilateral axons initially project medially along the lateral edge of the cardinal veins, in a similar path as the muscle progenitors, towards the nascent PPFs (Fig. 5). By E12.0 the phrenic axon is thickened and just reaches the nascent PPFs, which completely surround the distal tips of the axons (Fig. 5). Analysis of left and right phrenic nerves (Fig. 2D and data not shown) reveals that the right phrenic nerve axon consistently reaches the PPFs prior to the left phrenic nerve. Subsequently, the phrenic nerve axons elongate to accommodate the increasing distance between their roots in the cervical neural tube and their extension to the PPFs of the developing diaphragm.

In summary, our whole mount analysis of early embryos reveals the precise timing and spatial relationship of muscle migration and axon outgrowth of the phrenic nerve. The diaphragm’s muscle progenitors derive largely from somites C3 - C5 (with likely contribution from C2 and C6 somites) and during a 48 hour time window migrate initially in close proximity to the forelimb muscle progenitors, but then travel medially to the PPFs of the developing diaphragm. Starting significantly after the onset of muscle migration, the phrenic nerve axons extend towards the PPFs on a similar ventromedial pathway as the muscle progenitors. Interestingly, the right phrenic axon consistently reaches the PPFs prior to the left phrenic axon, but no consistent differences exist in the migration of left and right populations of muscle progenitors.

### Nascent PPFs are a medial extension of the lateral plate mesoderm

The PPFs are the target for the diaphragm’s muscle progenitors and axonal outgrowth of the phrenic nerve. As muscle migration and axonal outgrowth occurs between E9.5 and E11.5, this suggests that nascent PPFs are present during this early time period. Our previous studies have demonstrated the presence of PPFs by E11.5 (Merrell et al., 2015), but we did not examine their earlier development. We have shown that the *Prx1Cre* transgene labels the PPFs and their derivatives from E11.5 through P0 (Merrell et al., 2015). To investigate earlier PPF development we examined *Prx1Cre^Tg/+^;Rosa26^mTmG^* mice (more formally known as Prrx1Cre; Logan et al., 2002; Muzumdar et al., 2007) in whole mount at E10.0. During this time, no distinct, separate PPFs are present. Instead, consistent with detailed analysis by Durland and colleagues (2008), *Prx1Cre* mice label all the postcranial somatic lateral plate mesoderm (Fig. 4). The region that will become the PPFs is continuous with the entire lateral plate mesoderm that extends medially along the body wall (Fig. 4). Between E10.5 and E11.5 the nascent PPFs thicken and expand medially along the surface of the liver to become distinct pyramidal structures at E11.5 (Figure 5 and 6: Babiuk et al., 2003; Merrell et al., 2015).

**Figure 6:**
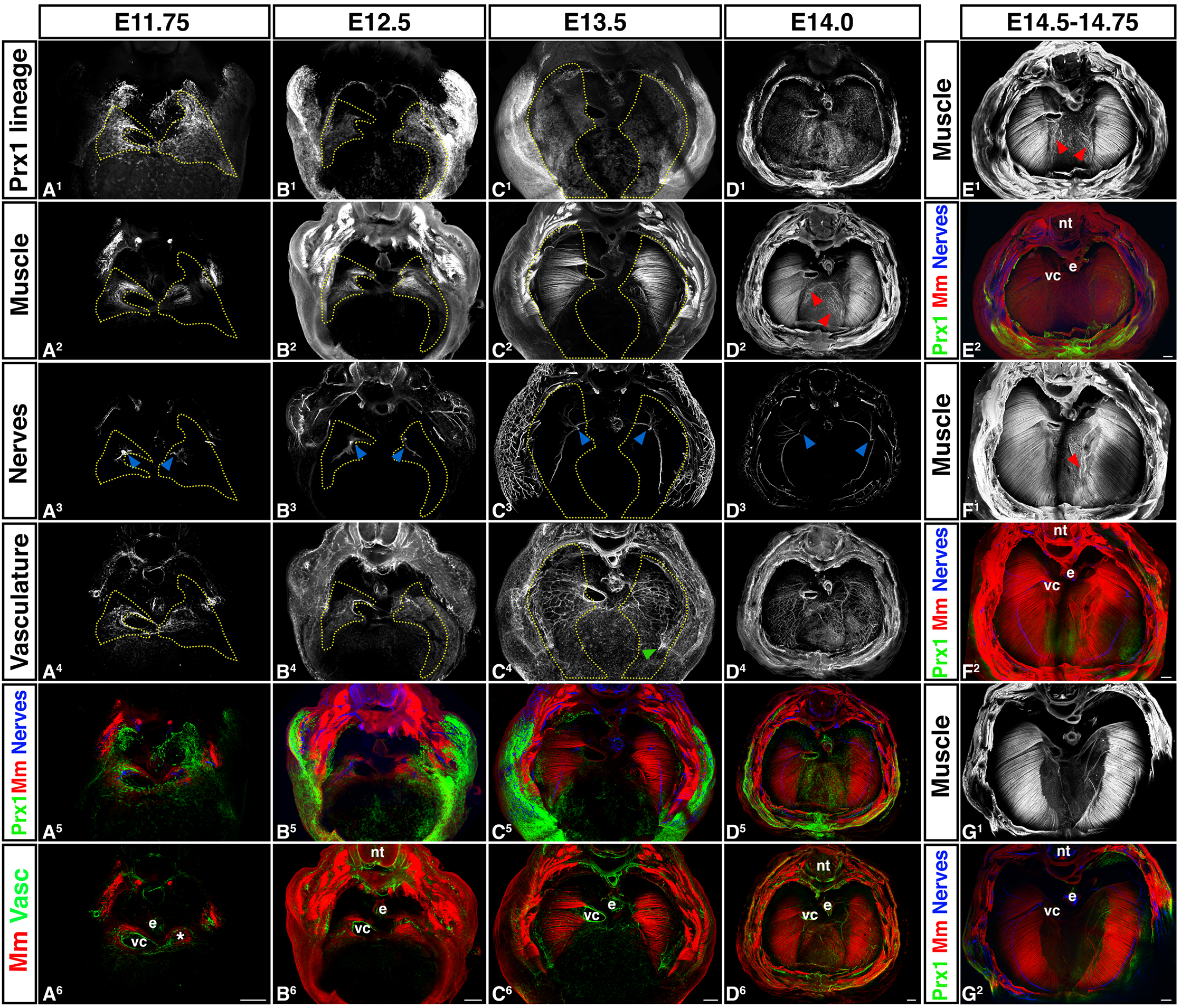
Morphogenesis of PPF-derived muscle connective tissue and central tendon, muscle, phrenic nerve, and vasculature are tightly linked temporally and spatially. (A) Muscle progenitors (A^2^) and axons of the phrenic nerve (A^3^) migrate into the right PPF (A^1^) adjacent to the vena cava (vc) and into the left PPF and caudal to the vestigial left horn of the sinus venosus (*). Vasculature (A^4^) also forms in PPF region in close association with vena cava and left sinus venosus horn. (B-D) PPFs migrate dorsally and ventrally to cover the surface of the underlying liver. In close association with the PPFs, muscle and vasculature expands as well as phrenic nerve axons extend dorsally and ventrally. Vessels from the body wall merge with the diaphragm’s vasculature (green arrow, C^4^). (D-G) Muscle progenitors appear and differentiate into myofibers in the central tendon region (D-F), but are gradually removed (G). nt, neural tube; e, esophagus; vc, vena cava. Scale bars A - G = 200um.

### Morphogenesis of PPF-derived muscle connective tissue and central tendon and muscle, are tightly linked temporally and spatially

Morphogenesis of the diaphragm begins with the appearance of distinct PPFs. In wholemount analysis of E11.75-12.0 *Prx1Cre^Tg/+^;Rosa26^mTmG^* embryos (Fig. 5 and 6A), distinct PPFs are present as pyramidal structures lying just cranial to the liver and septum transversum and the target of the muscle progenitors and the axons of the phrenic nerve. The PPFs have a complex geometry: dorsally, they extend to the cardinal veins and border the lung buds; laterally, they abut the body wall; and medially, the right PPF is lateral to the vena cava (the vein which connects to the right horn of the sinus venosus) and the left PPF is just caudal to the residual left horn of the sinus venosus. Of note, this complex geometry of the early PPFs has resulted in differing terms in previous studies, in which structures are not genetically labeled and are generally analyzed via sections. The PPFs, which we defined as the *Prx1*-derived pyramidal tissue sitting cranial to the septum transversum and liver (and target of muscle migration and phrenic nerve outgrowth; see Fig. S2 for delineation of PPFs in section), are equivalent to the PPFs of others (Babiuk et al., 2003; Clugston and Greer, 2007; Paris et al., 2015), but have also been referred to as the pleuroperitoneal membrane (Goodrich, 1958; Mall, 1910; Wells, 1954) or the post-hepatic mesenchymal plates (Carmona et al., 2016; Iritani, 1984; Kluth et al., 1987; Mayer et al., 2011). Also, the PPFs as defined by some studies (Carmona et al., 2016; Mayer et al., 2011) is equivalent to the region we define as the renal ridge, the caudal extension of the PPFs that extends to the developing kidneys and is devoid of muscle and phrenic nerve innervation (Fig. S2 E-F).

The PPFs (labeled via *Prx1Cre^Tg/+^;Rosa26^mTmG^*) expand dorsally and ventrally to cover the surface of the liver and give rise the central tendon and the muscle connective tissue associated with the costal diaphragm (Fig. 6 and Merrell et al., 2015). This expansion of the PPFs is independent of the presence of muscle (Merrell et al., 2015), but critically regulates development of the costal diaphragm’s muscle. Beginning at E11.75 muscle progenitors aggregate within the PPFs in close proximity to the vena cava on the right side or caudal to the residual left horn of the sinus venosus on the lefts side (Fig. 6A). As the PPFs expand, muscle progenitors are carried dorsally and ventrally and differentiate within the PPF-derived cells, but behind the leading edge of the PPFs (Fig. 6B-G; Merrell et al., 2015). By E15.0, congruent with the completed expansion of the PPF-derived cells across the liver’s surface, the morphogenesis of the costal muscle as a radial array of differentiated myofibers is nearly complete (Fig. 6D-G).

In the adult diaphragm the costal muscle is restricted to a peripheral ring of myofibers with no muscle present in the central tendon. Interestingly, via whole mount immunofluorescence (which is more sensitive than the methods employed in Merrell et al., 2015) we consistently observed (6/6 embryos at E14-14.75) ectopic muscle progenitors and myofibers in the central tendon region (Fig. 6D-G). These ectopic myofibers were generally aligned similarly with nearby costal myofibers and strongly vascularized, but not innervated (3 of 4 examined). We found in more mature diaphragms (as gaged by the ventral expansion of the muscle, Fig.6G) that little ectopic muscle is present, and in adult diaphragms muscle is completely absent from the central tendon region (data not shown). This suggests that during normal diaphragm development some muscle develops in the central tendon, but is selectively removed during diaphragm maturation.

### Phrenic nerve outgrowth and diaphragm vascularization are tightly linked temporally and spatially with costal diaphragm muscle morphogenesis

Correct targeting of phrenic nerve axons to the diaphragm’s muscle is critical for development of a functional diaphragm. The right phrenic nerve axon reaches the right PPF just lateral to the vena cava and the left phrenic nerve enters the left PPF at its medial side (Fig. 5 and 6A). By E11.75 (Fig. 6A), the phrenic nerve splits into three branches and by E12.5 the sternocostal branch projects ventrally, the dorsocostal branch projects dorsally, and the crural branch projects medially towards the crural muscle (Fig 6B). Subsequently, each of these branches elongates in close association with the dorsal and central expansion of the muscle (consistent with findings of Allan and Greer, 1997; Babiuk et al., 2003) and behind the leading of the PPFs (Fig 6C and D). Interestingly, while the initial projection of the right phrenic axon to the PPFs consistently precedes that of the left phrenic axon, the splitting and extension of the three branches does not continue to be more advanced than the left. Beginning by E13.5 the three primary branches extend secondary axons (Fig. 6C) and the pattern of primary and secondary branches is largely complete by E14.5 (Fig. 6. E-G). As has been noted by others (Charoy et al., 2017), the right phrenic nerve has more secondary branching and the angle between the dorsocostal and sternocostal branches is more acute than on the left side (Fig. 6D).

Vascularization of the diaphragm by the arterial network of the left and right phrenic, internal thoracic, and intercostal arteries is essential for diaphragm development and function (Stuelsatz et al., 2012). Using CD31 immunofluorescence to label the vasculature in whole mount, we found a small, but dense network of vessels is present in the PPFs and strongly associated with the muscle progenitors by E12.0 (Fig. 5 and 6A). As the muscle extends dorsally and ventrally, the vascular network expands in close association with the muscle (Fig. 6B-D). The vascular network anastomoses with vessels emerging from body wall musculature (Fig. 6C^4^). Distinct phrenic vessels descending along the dorsomedial edge of the costal diaphragm are visible by E14.0 (Fig. 6D^4^, D^6^). The internal thoracic arteries are not visible within the costal diaphragm at E14.0 and presumably appear at a lateral developmental time point.

The developmental origin of the diaphragm’s vasculature is unclear, and potentially migratory somitic cells or lateral plate cells could contribute endothelial cells. To determine whether somitic cells contribute to the endothelium we examined *Pax3^Cre/+^;Rosa26^mTmG^* embryos double labeled in whole mount via CD31 immunofluorescence and found a small number of GFP+ endothelial cells (Fig. S3A-D). To test whether lateral plate cells contribute to the endothelium, we examined *Prx1Cre^Tg/+^;Rosa26^mTmG^* embryos (which label the somatic lateral plate mesoderm) double labeled in whole mount via CD31 immunofluorescence, but found no GFP+ endothelial cells (Fig. S3E-H). Therfore, the somatic lateral plate does not contribute to the diaphragm’s endothelial and, similar to the limb (Hutcheson et al., 2009), a small number of endothelial cells derive from somitic migratory cells. What embryonic tissue is the source of the majority of the endothelial cells is still unclear, but likely the splanchnic portion of the lateral plate mesoderm (which is unlabeled by *Prx1Cre*; Durland et al., 2008).

## Discussion

Formation of the adult diaphragm requires the coordinated development of somite-derived muscle, neural tube-derived phrenic nerve, PPF-derived muscle connective tissue and central tendon, and vasculature. Diaphragm development is complicated due to the contribution of these multiple populations; the three-dimensional complexity of the nascent diaphragm and close association with heart, lungs, liver, and major vessels; and its caudal transposition during development. We have used wholemount immunofluorescence in combination with lineage tracing and three-dimensional reconstructions of mouse embryos to comprehensively analyze diaphragm development. Here we show that the key early developmental events (Fig. 7A) are the migration of muscle progenitors and projection of phrenic axons to the developing PPFs, whose development are tightly bounded by adjacent lungs, veins returning blood to the heart, and liver. Later morphogenesis of the diaphragm (Fig. 7B) involves PPF dorsal and ventral expansion and their differentiation into the central tendon and muscle connective tissue associated with the costal diaphragm muscle. Morphogenesis of the PPFs is tightly linked spatially and temporally with morphogenesis of the costal muscle, axonal outgrowth of the phrenic nerves, and formation of the vascular network. Therefore, the PPFs appear to be crucial for recruitment of muscle progenitors, targeting of phrenic axonal projections, and subsequent morphogenesis of the diaphragm.

**Figure 7:**
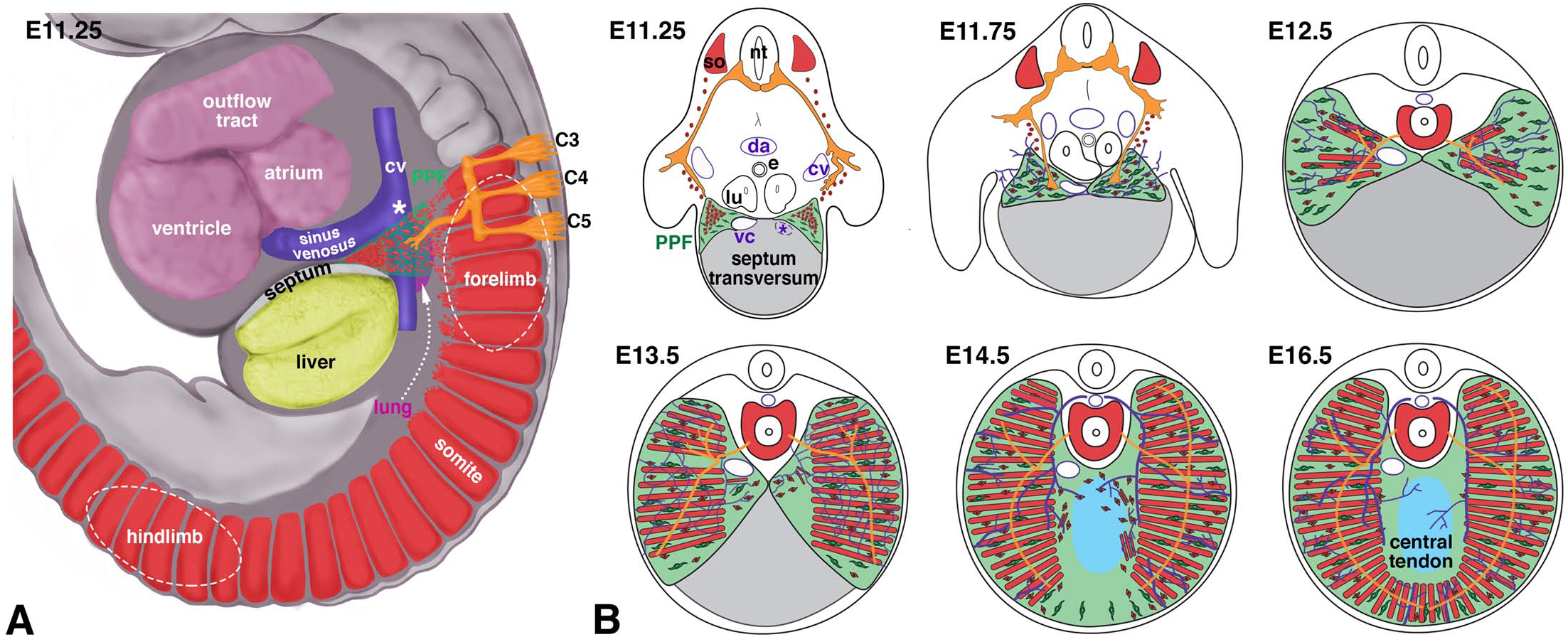
Model of diaphragm development. (A) At E9.5-11.5 muscle progenitors and phrenic nerve axons emigrate from C3-C5 region into nascent PPFs located between lung, sinus venosus, septum, and liver. (B) By E11.5 Muscle progenitors and phrenic nerve axons have targeted PPFs in vicinity of vena cava and caudal to left horn of sinus venosus. At E11.5-14.5 PPFs spread ventrally and dorsally and give rise to muscle connective and central tendon. Muscle, phrenic nerve, and vessels spread behind leading edge of PPFs. Some muscle initially differentiates in central tendon region. By E16.5 diaphragm morphogenesis is complete. cv, cardinal vein; vc, vena cava; sv, sinus venosus; so somite; da, dorsal aorta; e, esophagus; lu, lung; *, region just caudal to transient left horn of sinus venosus. Muscle (somites), red; PPFs, green; phrenic nerve, orange; vessels, purple.

Our whole mount analysis of a series of genetically and immunofluorescently labeled embryos reveals that the earliest event of diaphragm development is the migration of muscle progenitors from the somites to the nascent PPFs (Fig. 7A). Although previous analysis of sections of mouse embryos suggested that the diaphragm’s muscle originates from cervical somites (Dietrich et al., 1999), we explicitly demonstrate for the first time that the diaphragm’s muscle derives from a stream of muscle progenitors that emigrate from somites C3-5 (and potentially C2 and C6) beginning at E9.5. These muscle progenitors migrate just medial to the stream of forelimb muscle progenitors (that originate from somites C4-T1) and target the nascent PPFs. By E11.5 all diaphragm muscle progenitors have aggregated in the PPFs, either adjacent to the vena cava (in the right PPF) or caudal to the left horn of the sinus venosus (in the left PPF).

Beginning at E11.25 the axons of the phrenic nerve commence their projection towards the PPFs and follow a similar trajectory as the muscle progenitors. Thus we show, unlike previous reports based on sections (Jinguji and Takisawa, 1983)), that the initial muscle migration precedes projection of phrenic nerve axons and also occurs in a narrow window of time. After muscle progenitors and phrenic nerve axons reach the nascent PPFs, we find (consistent with observations of Allan and Greer, 1997) that the PPFs as well as the adjacent heart and associated vessels, lungs, and liver translocate caudally.

The target of muscle progenitor migration and the phrenic nerve axonal projection is the nascent PPFs. The PPFs develop as a medial outgrowth of the lateral plate mesoderm into the region bounded by the lungs, veins returning blood to the heart, and liver. The close proximity of the PPFs to the sinus venosus and the septum transversum covering the liver suggests that these two structures are important sources of signals directing the development of the PPFs. The septum transversum has been notably difficult to study because of the lack of markers or genetic reagents uniquely marking this structure and so its significance in early diaphragm development has not been definitively tested. The PPFs, in turn, are likely important sources of signals directing the migration of muscle progenitors and the projection of phrenic nerve axons. The identity of these signals and whether identical signals direct muscle and nerve are unknown, but an important avenue for future research.

The dorsal-ventral expansion of the PPFs is critical for diaphragm morphogenesis (Fig. 7B). As we show here and in a previous study (Merrell et al., 2015), between E12.5 and E14.5 the PPFs spread across the surface of the septum transversum and the liver. This expansion of the PPFs likely drives overall morphogenesis of the diaphragm; not only does the leading edge of the PPFs extend beyond that of the other tissues of the diaphragm, but this expansion is independent of muscle (Merrell et al., 2015). Dorsal-ventral expansion of muscle and vasculature closely track with one another and are slightly behind the leading edge of the PPFs. Consistent with the findings of others (Allan and Greer, 1997; Babiuk et al., 2003), the distal-most outgrowth of the sternocostal and dorsocostal branches of the phrenic nerve lies just behind the distal-most extent of the muscle. Therefore, the PPFs expand in advance of muscle and vasculature, which in turn are in advance of phrenic axon outgrowth. This suggests either that the PPFs directly control the morphogenesis of muscle, vessels, and nerve or that the PPFs control morphogenesis of muscle and vessels, which in turn relay morphogenetic signals to the nerve.

The PPF expansion gives rise to two domains: a lateral domain that forms muscle connective tissue and supports the costal muscle and a central domain that forms the central tendon and is muscle-less. How these domains arise has been unclear. Surprisingly, we found (via sensitive whole mount immunofluorescence) that during the early morphogenesis of the diaphragm, the central tendon region initially contains muscle progenitors, myoblasts, and differentiated myofibers. These centrally located patches of muscle become vascularized, but disappear so that the central region contains few, if any, myofibers by E15. The mechanism(s) controlling the selective removal of muscle from the central region are unclear, but may involve innervation; only a few of these central muscle patches were innervated and so the lack of innervation may lead to muscle death. Recently, Abelson tyrosine protein kinase 2 (Abl2) has been found to negatively regulate the growth of diaphragm muscle (Lee and Burden, 2017), and so potentially Abl2 expression in the central tendon region may lead to the exclusion of muscle in this area.

Defects in diaphragm development lead to CDH, and likely very early defects are an important cause of this birth defect. Previous CDH studies in mouse have shown that defects in diaphragm development are present (Carmona et al., 2016; Merrell et al., 2015) or genes are required (Paris et al., 2015) prior to E12.5, even though overt herniation does not happen until E16.5 (Merrell et al., 2015). We show that this is the time when PPFs are just forming and the target of migrating muscle progenitors and phrenic nerve outgrowth. As most CDH-implicated genes that have been examined are expressed in non-muscle PPF cells (Ackerman et al., 2005; Carmona et al., 2016; Clugston et al., 2008; Merrell et al., 2015; Paris et al., 2015), this implicates early defects in the PPFs as a source of CDH. Defects in the formation of the PPFs may directly cause CDH. Alternatively, PPFs may form, but be unable to signal to muscle progenitors and guide their migration to and targeting of the PPFs. A likely scenario underlying the etiology of many CDH cases is that genetic mutations directly affect the development of the PPFs, this has a non-autonomous effect on the neighboring muscle, and subsequently weakened muscle allows abdominal contents to herniate into the thoracic cavity and impede lung development. During the critical E9.5-12.5 time window, our analysis shows that the heart and associated vessels and lungs are in close proximity to the nascent PPFs and migrating muscle progenitors. Therefore, defects in molecular signals that affect PPF development may also affect nearby heart and lungs and thereby contribute to the heart and lung defects associated with CDH.

The preponderance of left-sided hernias in CDH has long been noted (Pober, 2007), but the underlying cause of this is unknown. We examined whether inherit asymmetries in diaphragm development could contribute to the predominance of left hernias. We noted no consistent differences in the timing of muscle migration to the PPFs, but others have noted that in neonatal mice the left half of the costal diaphragm is smaller than the right half (Charoy et al., 2017).

However, we did observe consistent differences between the outgrowth of left and right phrenic nerve axons. Axons of the right phrenic nerve consistently projected and reached the PPFs in advance of the left one and, as has been described others (Charoy et al., 2017), the geometry and secondary branching patterns of the right and left phrenic nerve differs. Thus, our study and others show that there are differences in the normal development of the left versus right halves of the diaphragm, but there is no clear indication of how these differences translate to the preponderance of left CDH.

The muscularized diaphragm is a critical defining feature of mammals, and how this evolutionary novelty arose at the base of mammals during the Permian era is a major unanswered question (Perry et al., 2010). Examination of the developmental innovations present in mammals, but absent from birds and reptiles is one method to elucidate the evolutionary origin of the diaphragm. Recruitment of muscle to the diaphragm during development is a crucial event present in mammals, but absent from birds and reptiles (Hirasawa et al., 2015; Hirasawa and Kuratani, 2013). Here we demonstrate that the diaphragm’s muscle progenitors originate from cervical somites, migrate in close association with the forelimb’s muscle progenitors into the somatic lateral plate mesoderm, but then diverge and migrate medially to the PPFs, and ultimately form the diaphragm’s muscle. In birds and mammals, cranial cervical somites give rise to neck muscles (Huang et al., 2000) and caudal cervical somites and cranial thoracic somites give rise to the forelimb muscles (Beddington and Martin, 1989; Beresford, 1983). However, only in mammals do the cervical somites give rise to muscle that migrates to the diaphragm region. Therefore, key events in the development and evolution of the diaphragm are recruitment of muscle progenitors from the cervical somites and guidance of these progenitors to the nascent diaphragm. Determining the molecular and cellular mechanisms regulating these events will be key for elucidating the evolutionary origin of the diaphragm, a functionally critical and defining feature of mammals.

## Acknowledgements

We thank C. Rodesch and M. Redd at the University of Utah Imaging Core and Y. Wan and C. Hansen for help with Fluorender analysis. We thank E. Bogenschutz, B. C. Collins, and C. Shaw for critical reading of the manuscript. Fluorender is supported by NIH R01 EB023947 and P41 GM103545. E.M.S. is supported by NIH F32 HD093425. This research was supported by NIH R01 HD087360; March of Dimes FY15-203; and the Wheeler Foundation to G.K.

---

**Supplemental Table 1.** Antibodies used in study

| **Antibody** | **Type** | **Source** | **Product Number** | **Working Concentration** | **Secondary and Amplification** |
| --- | --- | --- | --- | --- | --- |
| Pax7 | Mouse IgG1 | DSHB | PAX7 | 2.4 µg/ml | Alexa 647 goat α mouse IgG1 |
| MyoD | Mouse IgG1 | Thermo Fisher | MA5-12902 | 4 µg/ml | Alexa 647 goat α mouse IgG1 |
| MyHC (fast, neonatal) My32 | Mouse IgG1 | Sigma | M4276 | 10 µg/ml | Alexa 647 goat α mouse IgG1 |
| GFP | Chick polyclonal | Aves Labs | GFP-1020 | 20 µg/ml | Cy2 goat α chick |
| Neurofilament-L | Rabbit monoclonal | Cell Signaling | 2837 | 0.48 ug/ml | Alexa 594 goat α rabbit or Alexa 405 goat α rabbit or HRP goat α rabbit |
| Neurofilament | Mouse IgG1 | DSHB | 2H3 | 2 µg/ml | Alexa 647 goat α mouse IgG1 |
| CD31 | Rat | BD Pharmingen | 557355 | 2.5 µg/ml | Biotin goat α rat, Cy3 Conjugated Streptavidin |
| Tomato | Rabbit polyclonal | Clontech | 632496 | 1:200 | Alexa 594 goat α rabbit |

**Figure S1:**
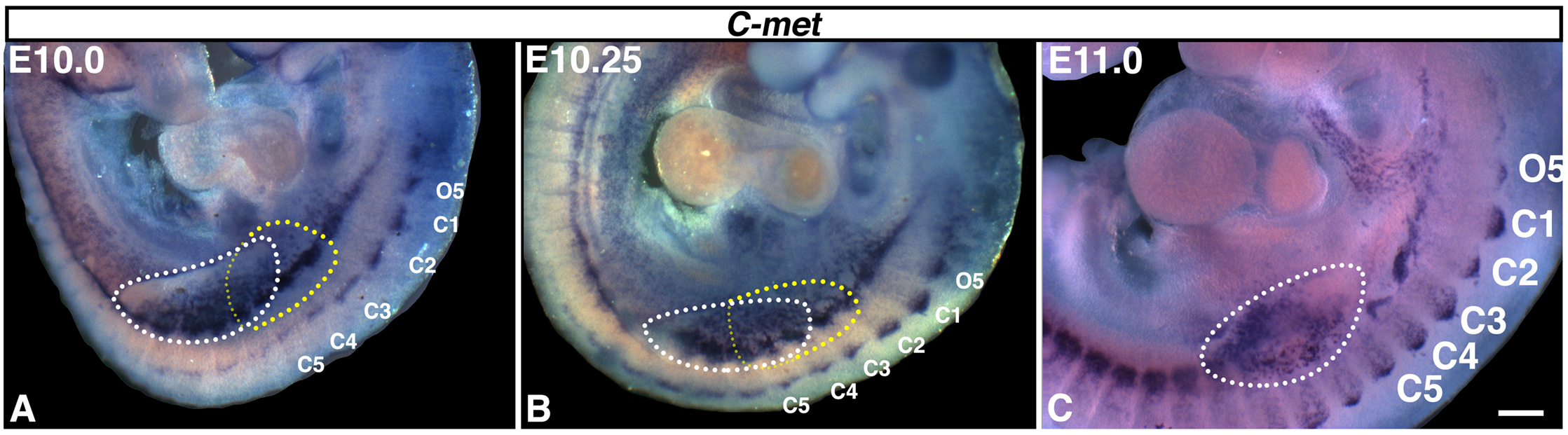
*Met*+ diaphragm progenitors delaminate from C3-5 somites. *Met* in situ hybridization shows delamination of diaphragm muscle progenitors (outlined in yellow) in close proximity to limb muscle progenitors (outlined in white). By E11.0 most diaphragm progenitors are medial to limb muscle. Scale bar A-C = 200um.

**Figure S2:**
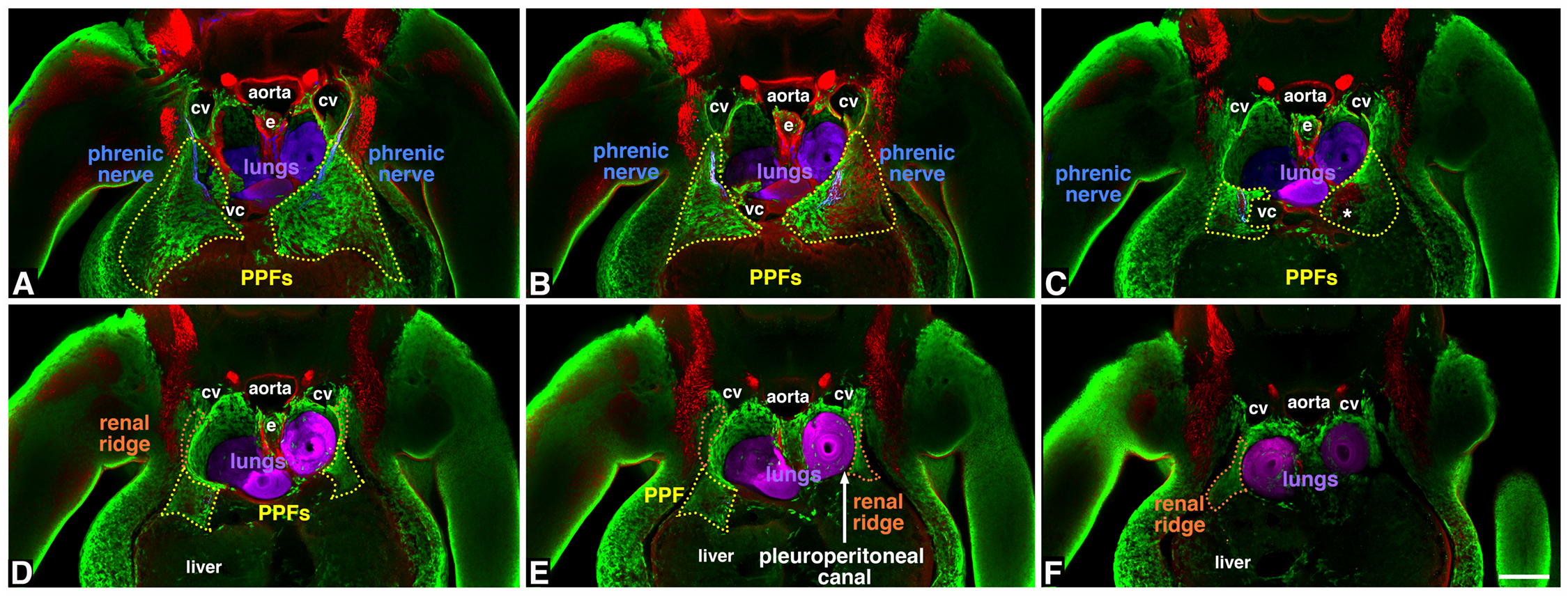
Optical serial sections show complex geometry of PPFs. (A-D) Cranial to caudal sections through E12.0 *Prx1Cre^Tg/+^;Rosa26^mTmG/+^* diaphragm region. (A-C) Cranial-most sections of PPFs (green) contain muscle progenitors and phrenic nerve axons. (D-E) More caudal regions of PPFs are devoid of muscle and nerve and lie dorsal to the liver and ventral to the lungs. (F) Renal ridge is the caudal extension of the PPFs. Scale bar A-F = 200um. Sections A-E are each 30 um apart, while section F is 60um caudal to section E. Sections are through embryo shown in Fig. 5.

**Figure S3:**
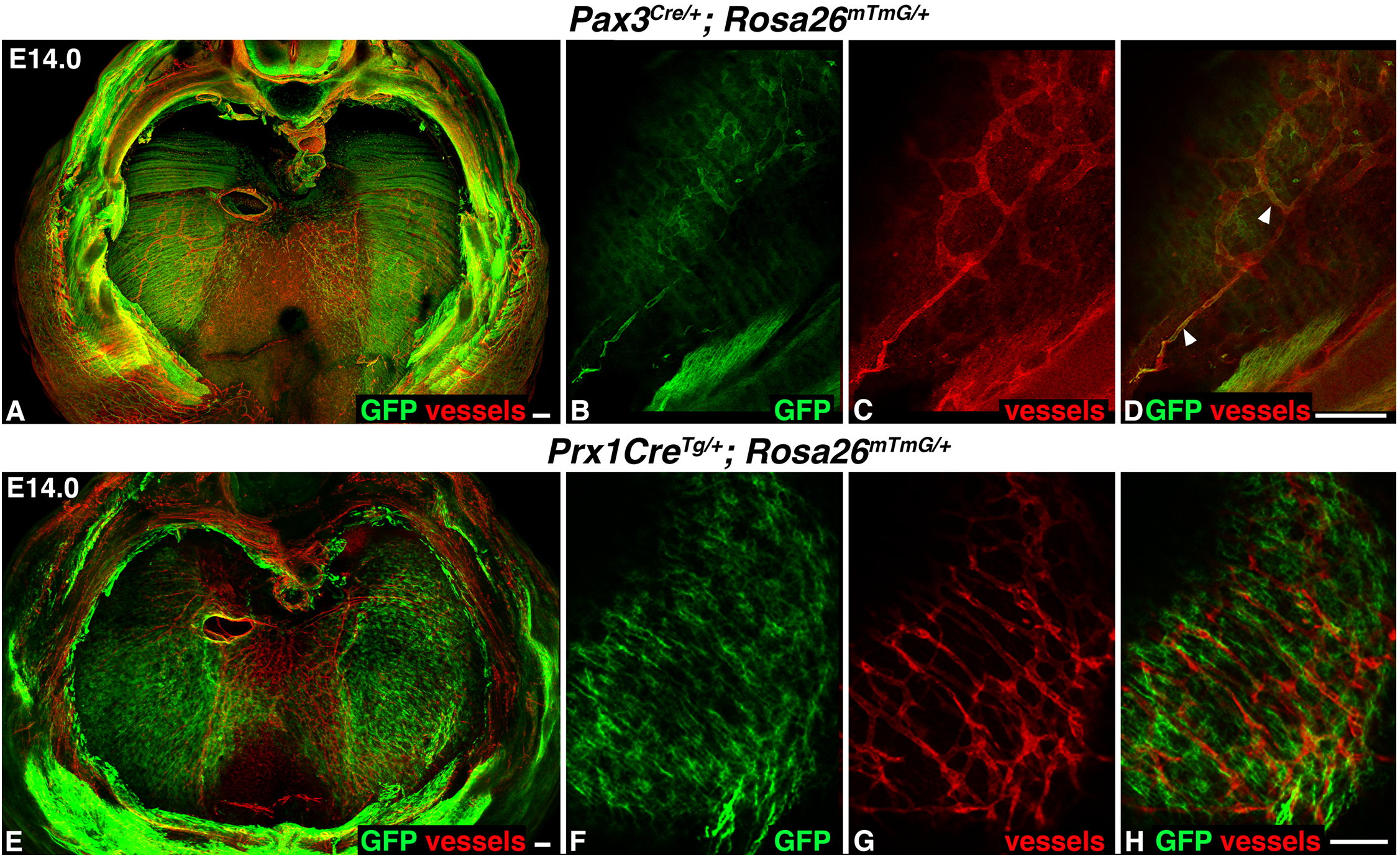
A few endothelial cells are derived from the somites. (A-D) In whole mount (A) or sections (B-D) somite derivatives labeled in green via *Pax3^/+^;Rosa26^mTmG/+^* give rise to a small number of red CD31+ immunofluorescently labeled endothelial cells of vessels. (E-H) In whole mount (E) or sections (F-H) PPF derivatives labeled in green via *Prx1Cre^Tg/+^;Rosa26^mTmG/+^* do not give rise to red CD31+ endothelial cells. cv, cardinal vein; vc, vena cava; e, esophagus; *, transient right horn of sinus venosus. Scale bar A, B-D, E, F-H = 100um.

